# Heritability and transcriptional impact of JAK3, STAT5A and STAT6 variants in a Tyrolean family

**DOI:** 10.64898/2025.12.14.694240

**Authors:** Hye Kyung Lee, Teemu Haikarainen, Yasmine Caf, Priscilla A. Furth, Ludwig Knabl, Olli Silvennoinen, Lothar Hennighausen

**Author notes:** Correspondence: Lothar Hennighausen and Hye Kyung Lee. Co-first author (Hye Kyung Lee, Teemu Haikarainen). co-last author (Olli Silvennoinen, Lothar Hennighausen).

## Abstract

The Janus Kinase (JAK) and Signal Transducers and Activators of Transcription (STAT) pathway regulate a range of biological processes, including immune response and hematopoiesis. While a major research focus has been on somatic human mutations in disease, less is known about the heritability of germline variants and their physiological impact. Here we identify the rare JAK3^P151R^, JAK3^R925S^, STAT5A^V494L^ and STAT6^Q633H^ variants in an extended family spanning three generations and integrate *in silico* analyses, AlphaFold 3 structural predictions and investigate the immune transcriptomes in probands carrying one or more variants. All four variants are inherited through the germline without any evident clinical or physiological manifestations in the carriers. As individual variants, not all persons carrying a specific variant showed the same immune transcriptome. The presence of activated basal transcriptomes was limited to some, but not all, individuals carrying the above variants above. A next step in understanding the role of germline variants will be to understand how and why other factors including both other germline variants as well as environmental and developmental factors influence the likelihood of expression of an activated basal transcriptome.

## Introduction

Janus Kinases (JAKs) and Signal Transducers and Activators of Transcription (STATs) play indispensable roles in translating cytokine signals into transcriptional responses that regulate a range of biological processes, including body growth^1^, lactation^2^, hematopoiesis^2-4^, inflammation and antiviral defense. Dysregulated JAK/STAT signaling has been implicated in chronic inflammation and autoimmune disease, and somatic variants have been identified predominantly in hematological malignancies. These include the classical JAK2^V617F^ variant in myeloproliferative neoplasms (MPNs^5,6^ and SH2 domain variants of STAT family members^7,8^.

We recently identified a range of germline JAK/STAT variants and linked some to an exacerbated innate immune response after vaccination^9,10^ and an elevated baseline immune transcriptome^11^. Access to community-based cohorts permitted us to address yet unanswered questions focused on the presence of JAK/STAT variants in large families. First, it is not well understood to what extent these variants are transmitted through the germline, especially in individuals carrying two or more variants. Second, the combinatorial transcriptional impact of several variants in individuals remains to be understood. Here we identified a large family carrying two JAK3 variants and one STAT5A and STAT6 variant. We used *in silico* analyses and AlphaFold 3 (AF3) protein structure predictions to assess the pathogenicity of these variants and gauged their impact in individuals carrying one or more variants.

## Material and Methods

### Study participant details

Identification of Single Nucleotide Variations (SNVs) was conducted on n=13 publicly available RNA-seq datasets derived from a study of a family infected with the SARS-CoV-2 Omicron strain^12,13^. WES was conducted on nine of these individuals and an additional six members of this family. Thus, SNV data were obtained from a total of 17 family members (Figure 1). N=9 females (age 49 +/-17 years) and n=8 males (age 45 +/-15 years) from Tyrol (Austria)^12^. All participants in the study were of Austrian ethnicity. The influence of sex, gender, or both cannot be reported from this study due to the small group size. This is a limitation of the research’s generalizability. The Institutional Review Board (IRB) of the Office of Research Oversight/Regulatory Affairs, Medical University of Innsbruck, Austria, which is responsible for all human research studies conducted in the State of Tyrol (Austria) approved the study (EK Nr: 1064/2021). All individuals provided written informed consent, all information obtained was coded and anonymized, and all methods were carried out in accordance with relevant guidelines and regulations. The research was performed in accordance with the Declaration of Helsinki (https://www.wma.net/policies-post/wma-declaration-of-helsinki-ethicalprinciples-for-medical-research-involving-human-subjects/).

**Figure 1.**
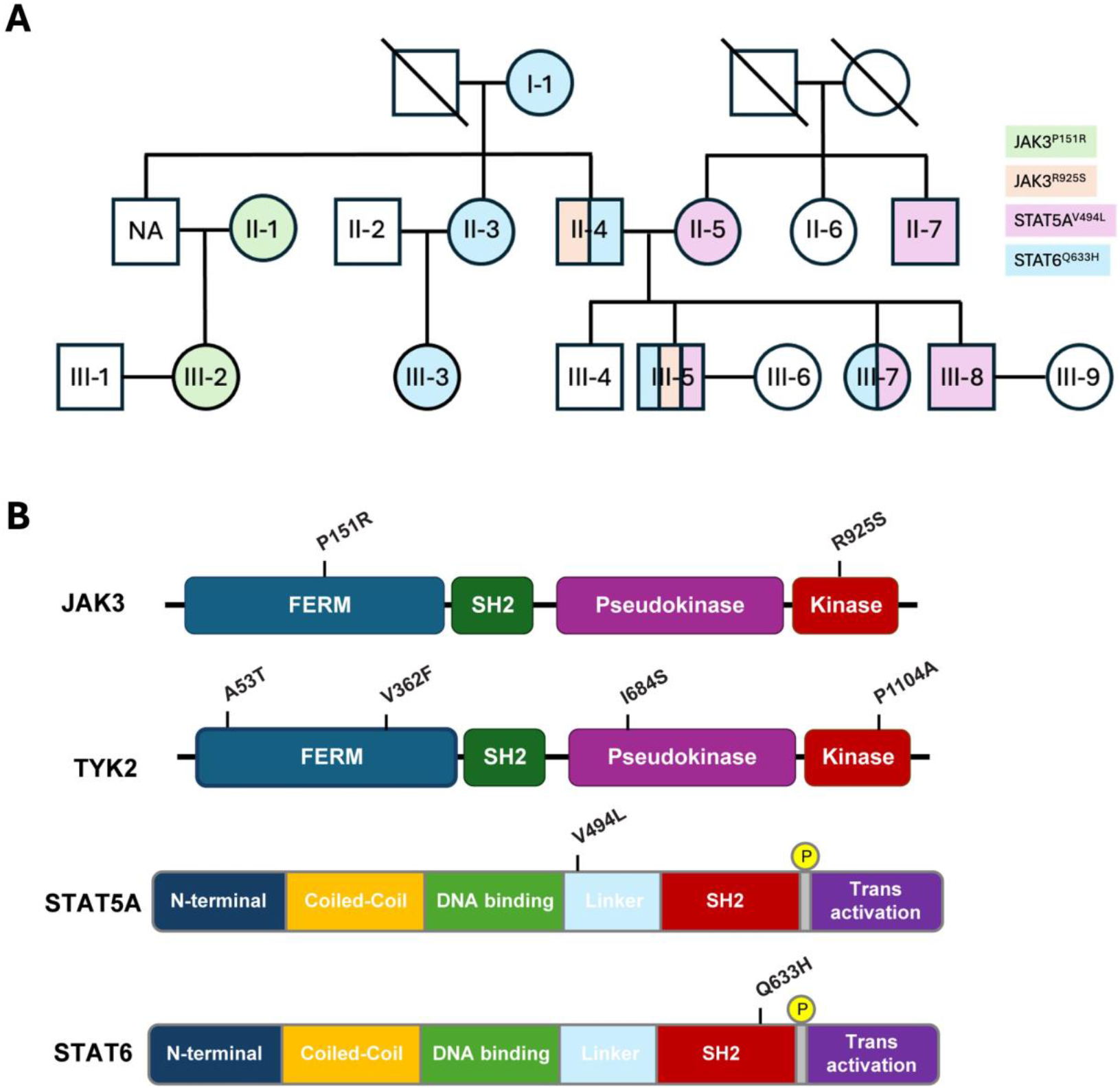
Identifcation of JAK/STAT variants in an extended Tyrolean family. **(A)** Family tree. **(B)** Domain structures of JAK3, STAT5A and STAT6 and location of the identified variants. JAK3^P151R^ is located in the FERM domain, JAK3^R925S^ in the kinase domain, STAT5A^V494L^ in the linker between the DBD and the SH2 domain. STAT6^Q633H^ is located in the SH2 domain.

### Whole genome sequencing and data analysis

Genomic DNA was extracted from human buffy coat samples using the Wizard Genomic DNA Purification Kit (Promega). PCR-free library preparation and whole-genome sequencing were performed at the NIH Intramural Sequencing Center (NISC). In this process, PCR-free libraries were constructed from 1μg of genomic DNA utilizing the TruSeq DNA PCR-Free HT Sample Preparation Kit (Illumina). The median fragment size of the libraries was approximately 400 base pairs (bp). Each library was tagged with unique dual-index DNA barcodes to facilitate library pooling while minimizing the occurrence of barcode hopping. Libraries were pooled and sequenced on the NovaSeq 6000 or NovaSeq X + (Illumina) platforms, generating a minimum of 300 million paired-end reads (151 base pairs each) per individual library.

Subsequent data analysis was conducted using the Clara Parabricks Pipeline (version 4.0.1) for the variants’ detection (https://docs.nvidia.com/clara/parabricks/4.0.1/whatsnew.html). Alignment of the sequencing reads was performed with BWA MEM^14^ using the human reference genome (hg38), followed by duplicate marking with Picard tools^15^. The Genome Analysis Toolkit (GATK) pipeline was employed for base quality score recalibration (BQSR) and variant calling, with HaplotypeCaller generating gvcf files using a minimum Phred-scaled confidence threshold of 16.

### Single Nucleotide Variant (SNV) detection in RNA-seq data

The raw data generated from RNA of human buffy coat were subjected to QC analyses using the FastQC tool (version 0.11.9) (https://www.bioinformatics.babraham.ac.uk/projects/fastqc/). mRNA-seq read quality control was done using Trimmomatic^16^ (version 0.36) and STAR 2-pass procedure^17^ (version STAR 2.7.9a) was used to align the reads (hg19). Aligned reads were filtered using BWA MEM^14^ (version 0.7.15), followed by Picard tools^15^ (version 2.9.2) to mark duplicates. The GATK analysis workflow was applied as follows: (i) base recalibration - GATK BaseRecalibrator, AnalyzeCovariates, and PrintReads - using the databases of known polymorphic sites, dbSNP138 (provided by the high-performance computing team of the NIH (Biowulf)); (ii) variant calling - GATK HaplotypeCaller - with the genotyping mode “discovery”, the “ERC” parameter for creating gvcf and a minimum phred-scaled confidence threshold of 30. Hard filters were applied: QD < 2.0 || QUAL < 30.0 || SOR > 3.0 || FS > 60.0 || MQ < 40.0 || MQRankSum < -12.5 || ReadPosRankSum < -8.0. The resulting SNVs were additionally filtered by removing those overlapping with repetitive elements^18^ (UCSC’s masked repeats plus simple repeats; https://hgdownload.soe.ucsc.edu/goldenPath/hg19/database/) and black regions (ENCODE^19^; https://mitra.stanford.edu/kundaje/akundaje/release/blacklists/hg19-human/). On an individual level, only SNVs with a genotype of 0/1 or 1/1 were kept. Further filtering steps comprised the removal of SNVs with a read depth smaller than 10, an excessive read depth^20^ (d+3√d, d = average read depth), an allele frequency less than 10% using a variety of tools (BEDtools, version 2.26.0; BEDOPS, version 2.4.3; VCFtools, version 0.1.17)^21-23^. All SNVs within +/-5bp of an indel border were removed as likely false-positives. SNP position on hg19 was converted to hg38.

### mRNA sequencing (mRNA-seq) data analysis

Ribosomal RNA was removed from 1 μg of total RNAs and cDNA was synthesized using SuperScript III (Invitrogen). Libraries for sequencing were prepared according to the manufacturer’s instructions with TruSeq Stranded Total RNA Library Prep Kit with Ribo-Zero Gold (Illumina, RS-122-2301) and paired-end sequencing was done with a NovaSeq 6000 instrument (Illumina).

The raw data were subjected to QC analyses using the FastQC tool (version 0.11.9). mRNA-seq read quality control was done using Trimmomatic^16^ (version 0.36) and STAR RNA-seq^17^ (version STAR 2.7.9a) using 150 bp paired-end mode was used to align the reads (hg38). HTSeq^24^ (version 0.9.1) was to retrieve the raw counts and subsequently, Bioconductor package DESeq2^25^ in R (https://www.R-project.org/) was used to normalize the counts across samples^26^ and perform differential expression gene analysis. Additionally, the RUVSeq^27^ package was applied to remove confounding factors. The data were pre-filtered keeping only genes with at least ten reads in total. The visualization was done using dplyr (https://CRAN.R-project.org/package=dplyr) and ggplot2^28^. P-values were calculated using a paired, two-side Wilcoxon test and adjusted p-value (pAdj) corrected using the Benjamini–Hochberg method. The cut-off value for the false discovery rate was pAdj > 0.05. Genes with log^2^ fold change >1 or <-1, pAdj <0.05 and without 0 value from all sample were considered significant and then conducted gene enrichment analysis (GSEA, https://www.gsea-msigdb.org/gsea/msigdb). For significance of each GSEA category, significantly regulated gene sets were evaluated with the Kolmogorov-Smirnov statistic. A value of \**P* < 0.05, \*\**P* < 0.01, \*\*\**P* < 0.001, \*\*\*\**P* < 0.0001 was considered statistically significant.

### Allele frequency information and *in silico* prediction of altered function

Allele frequency information for certain variants was collected using multiple databases, like dbSNP (https://www.ncbi.nlm.nih.gov/snp/), gnomAD (https://gnomAD.broadinstitute.org/), All of Us (https://workbench.researchallofus.org), COSMIC (https://cancer.sanger.ac.uk/cosmic) and ClinVar (https://www.ncbi.nlm.nih.gov/clinvar/).

Multiple computational tools were employed to assess the potential for altered function of the JAK/STAT variants. AlphaMissense^29^ (https://alphamissense.hegelab.org/search) was used to predict the functional impact of variants, with scores ranging from 0 (benign) to 1 (pathogenic). The Rare Exome Variant Ensemble Learner (REVEL)^30^ scores, which range from 0 to 1, were calculated to assess the probability of pathogenicity, as defined by the program. Additionally, PolyPhen-2^31^ (http://genetics.bwh.harvard.edu/pph2/bgi.shtml) analysis was performed to predict the possible impact of amino acid substitutions on protein structure and function, with scores ranging from 0 (benign) to 1 (probably damaging).

### Structural analyses

Monomeric JAK3 structure was predicted using AlphaFold3 (https://alphafoldserver.com). Structural analysis and visualization were performed using PyMOL (version 3.1.6.1).

### Data availability

All data were obtained or uploaded to Gene Expression Omnibus (GEO). The RNA-seq generated in this study are available using accession code GSE313264. The RNA-seq data of Omicron patients were obtained using accession number GSE201530^12^ and GSE205244^13^. The whole genome sequencing data generated in this study has not been deposited in a public database due to privacy concerns. Researchers interested in accessing the dataset may contact the authors directly.

## Results

In a quest to understand the heritability of JAK/STAT variants and further explore their combined transcriptional impact, we searched for families carrying two or more variants in any of the three JAK and seven STAT proteins. Here we report on a family we had recruited during the COVID-19 pandemic to investigate the innate immune response after COVID-19 Omicron infection^12^. Both RNA-seq and whole exome sequencing (WES) data collected from 17 members of a Tyrolean family spanning three generations revealed two JAK3 variants and one variant each in STAT5A and STAT6 were identified (Figure 1). The JAK3^P151R^ and JAK3^R925S^ variants are in the FERM domain and the kinase domain, respectively. The STAT5A^V494L^ variant is within the linker separating the DNA Binding Domain (DBD) and the SH2 domain, and the STAT6^Q633H^ variant is positioned within the SH2 domain. The STAT6^Q633H^ variant was identified in six individuals in three generations and the STAT5A^V494L^, JAK3^P151R^ and JAK3^R925S^ variants in two generations. Two individuals carried only a single variant, one person carried the STAT6^Q633H^ and JAK3^R925S^ variants, one the STAT6^Q633H^ and STAT5A^V494L^ variants and one person carried three variants, STAT6^Q633H^, STAT5A^V494L^ and JAK3^R925S^ (Supplementary Table 1). Both JAK3^P151R^ and JAK3^R925S^ are rare variants, STAT5A^V494L^ has not been listed in databases and STAT6^Q633H^ is an ultra-rare variant with an allele frequency of approximately 10^-4^ in GnomAd and AllofUs (Table 1). Most individuals in this family also carried TYK2 mutations (Supplementary Table 1). These findings demonstrate that all four variants are inherited through the germline without any evident clinical or physiological manifestations in the carriers. Importantly, individual II-4 passed on the STAT6^Q633H^ and JAK3^R925S^ variants to his son who also inherited the STAT5A^V494L^ variant from his mother, II-5 (Figure 1).

**Table 1.**
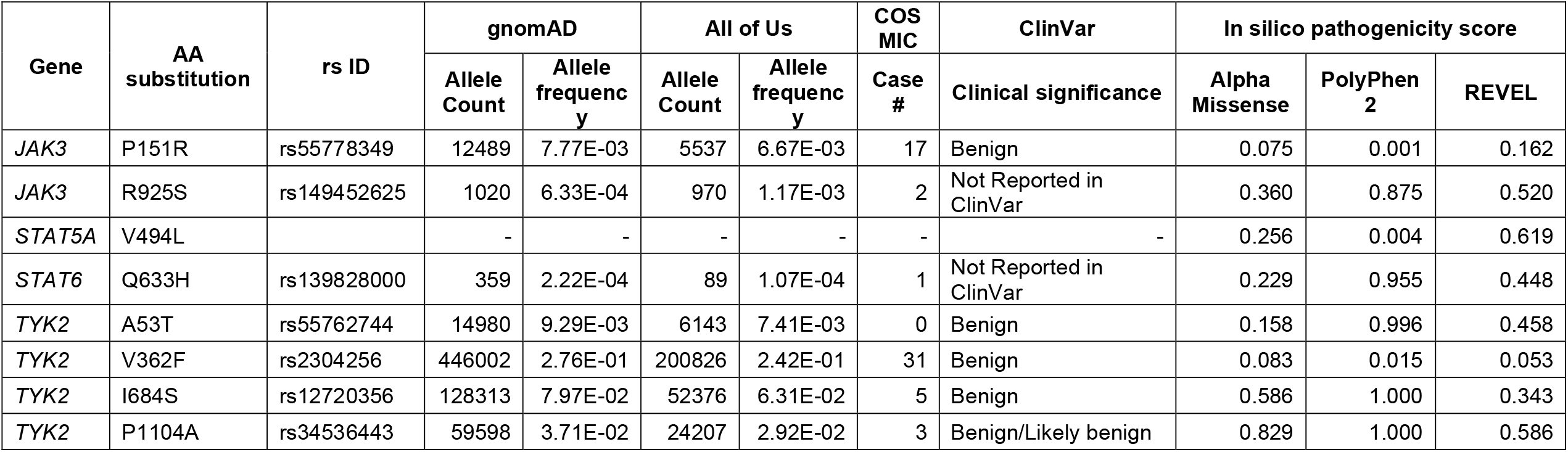
JAK and STAT variants identified in study subjects.

To evaluate the potential pathogenicity of these four variants, we employed the NIH ClinVar database and additional in silico prediction tools (Table 1). While no ClinVar records for were available for STAT6^Q633H^, STAT5A^V494L^ and JAK3^R925S^, JAK3^P151R^ was considered as benign. AlphaMissense^29^, a state-of-the-art computational tool, predicted an ambiguous score for JAK3^R925S^ and benign scores for the other three variants. The PolyPhen-2^32^ scores predicted a significant impact of the JAK3^R925S^ and STAT6^Q633H^ variants. The REVEL (Rare Exome Variant Ensemble Learner)^30^ scores indicated a higher probability of pathogenicity of JAK3^R925S^, STAT5A^V494L^ and STAT6^Q633H^.

## Structural analysis of JAK and STAT mutations

JAK3 has very little structural information available compared to other family members, and only the structure of JAK3 kinase domain has been experimentally resolved. Therefore, AI-based structure prediction with AF3 was used to obtain structural information of the two identified mutations. Notably, structure prediction of JAK3 with AF3 results in a highly similar tertiary structure when compared to JAK1, JAK2 and TYK2. We also analyzed the mutations in the context of a cryo-EM structure of activated JAK dimer (PDB code 8EWY) but in the activated conformation both mutated residues faced solvent and did not participate in intra-domain or JAK-JAK interaction across the dimer interface. Therefore, it is likely that the mutations exert their effects, if any, on the monomeric, autoinhibited conformation.

The P151R mutation is located in the FERM-SH2 domain in a short linker between two α-helices (Figure 2A). The residue is solvent exposed and based on the structural analysis, the mutation would not disrupt or create any specific interactions. However, due to its rigid cyclic structure, introducing a more flexible residue in the place of proline could lead to disruption of the linker and the conformation of the protein fold, at least locally. The R925S mutation is positioned in the C-lobe of the kinase domain and in the AF3 model the arginine side chain faces the SH2-JH2 linker which runs in the interface of FERM-SH2, pseudokinase, and kinase domains (Figure 2A, B). The mutation could disrupt the linker conformation and thus the autoinhibited conformation of JAK3. Importantly, the SH2-JH2 linker is a hotspot of pathogenic JAK mutation, and harbors activating mutations especially in hematologic malignancies, e.g. JAK3 M511I^33^ and JAK2 K539L^34^.

**Figure 2.**
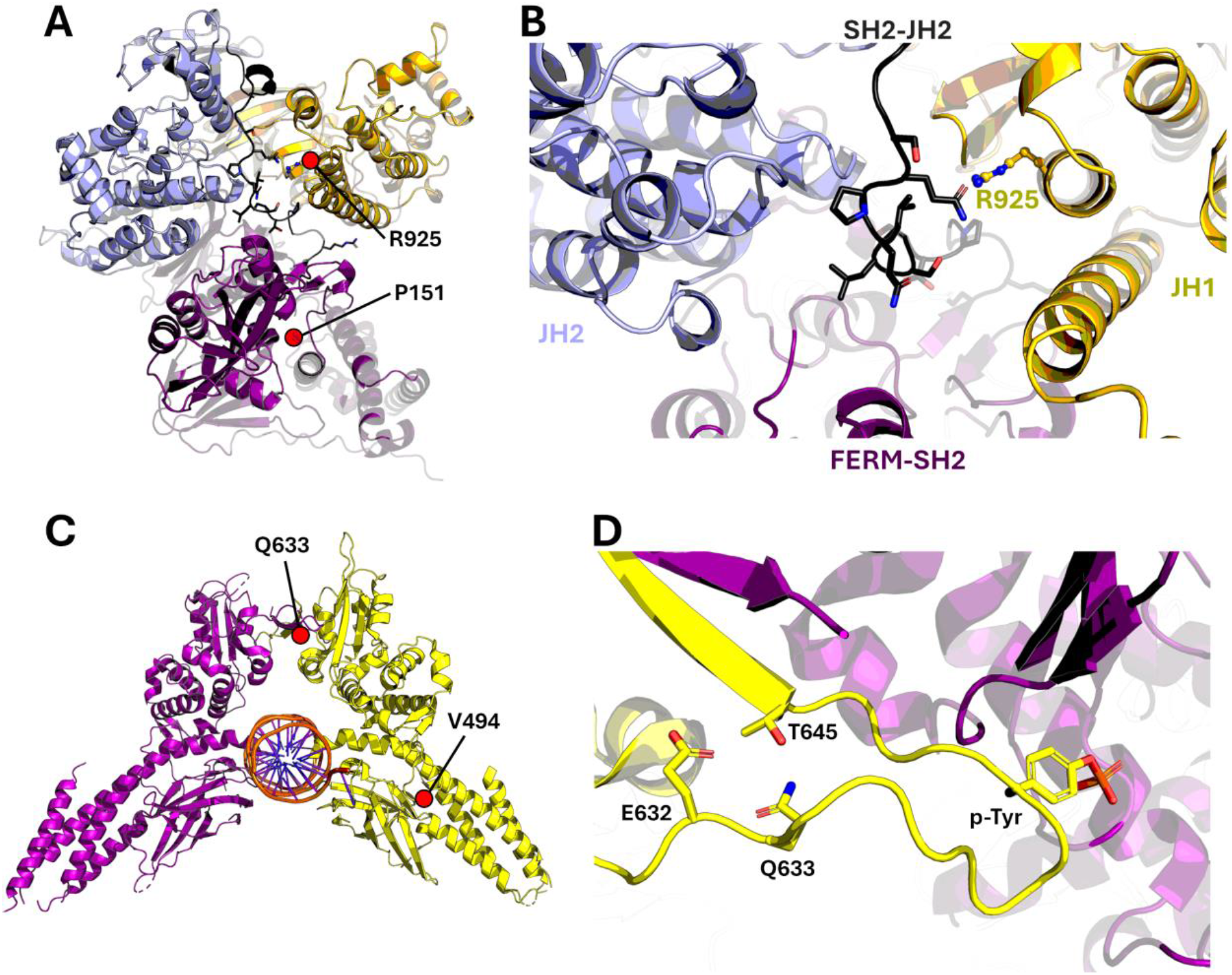
Structural analysis of the mutants. **(A)** The location of Pro125 and Arg925 in the AF3 model. **(B)** The arginine side chain points towards the SH2-JH2 linker. FERM-SH2 is colored in purple, JH2 in light blue, JH1 in yellow, and the SH2-JH2 linker in dark gray. **(C)** STAT6 structure (PDB code 4Y5W)^43^ showing the location of STAT5A V494L and STAT6 Q633H mutations. **(D)** Close-up view of the STAT6 Q633H mutation next to the phosphotyrosine (p-Tyr) binding site.

The identified STAT5A V494L mutation is located in a solvent exposed loop at the end of the DNA-binding domain (Figure 2C). Wild-type valine does not participate in any direct interactions via its side chain, which points towards the solvent. Therefore, the conservative Val to Leu mutation is likely benign. STAT6 Q633H is in the SH2 domain next to the phosphotyrosine binding site responsible for STAT dimerization (Figure 2C and 2D). The glutamine residue makes a hydrogen bond with Thr645 contributing to the stability of the phosphotyrosine-containing loop. The Q633H likely conserves this interaction as histidine can form similar hydrogen bond via its side chain. No drastic effects are therefore suspected from this mutation.

## STAT5A and STAT6 variants are associated with enhanced basal immune transcriptomes

To explore the potential impact of these JAK/STAT variants on the baseline immune transcriptome in a real-world setting, we conducted RNA-seq on PBMC from 13 family members (Supplementary Table 1). Specifically, we had baseline transcriptome data from five participants carrying the STAT6^Q633H^ variant. Three members (I-1, II-3 and III-3), spanning three generations carried only the STAT6^Q633H^ variant, while III-7 also carried the STAT5A^V494L^ variant. Individual III-5 carried both the STAT6^Q633H^ and the STAT5A^V494L^ variants as well as the JAK3^R925S^ variant. Volcano plots were generated to visualize differentially expressed genes (Figure 3). While a significant number of genes were deregulated in individual I-1 (STAT6^Q633H^ variant),, her daughter and granddaughter did not display an aberrant immune transcriptome. The differences observed between these three individuals could be explained be age differences, with I-1 being 87 years old, II-3 55 years and III-3 36 years old. Alternatively, additional mutations in immune regulatory pathways could explain the result. Individual III-5 carrying the STAT6^Q633H^ variant in combination with the STAT5A^V494L^ and JAK3^R925S^ variants also presented an elevated transcriptome. Among the individuals carrying the STAT5A^V494L^, II-5 did display an enhanced transcriptome. The presence of various combinations of TYK2 variants in these individuals (Suppl. Table 1) complicates the interpretation of the findings.

**Figure 3.**
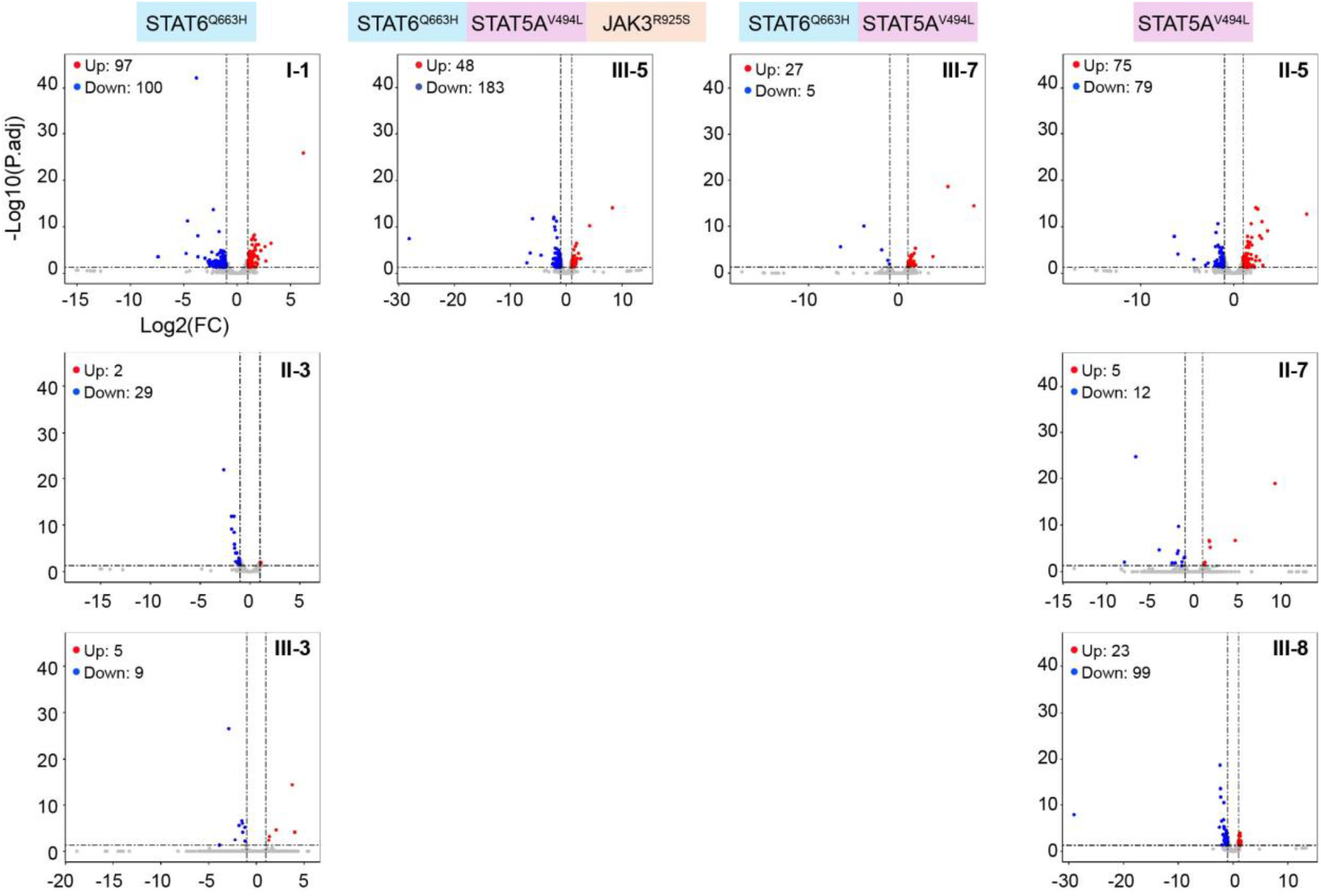
Differentially expressed genes in PBMCs from healthy probands carrying different combinations of JAK/STAT variants.

We had previously analyzed RNA-seq data from a cohort infected with the SARS-CoV-2 Omicron strain^12^, including 13 individuals from the family investigated in this study. Blood had been collected approximately two weeks after the onset of symptoms and RNA-seq was conducted on PBMCs. Volcano plots demonstrated the highest induction of gene expression in I-1 (STAT6^Q633H^), II-5 (STAT5A^V494L^), III-7 (STAT6^Q633H^; STAT5A^V494L^), III-5 (STAT6^Q633H^; STAT5A^V494L^; JAK3^R925S^) and II-4 (STAT6^Q633H^; JAK3^R925S^) (Figure 4). Notably, individuals with elevated transcriptomes after SARS-CoV-2 Omicron infection also had elevated baseline transcriptomes (Figure 3), suggesting a genetic component, possibly the JAK/STAT variants described here.

**Figure 4.**
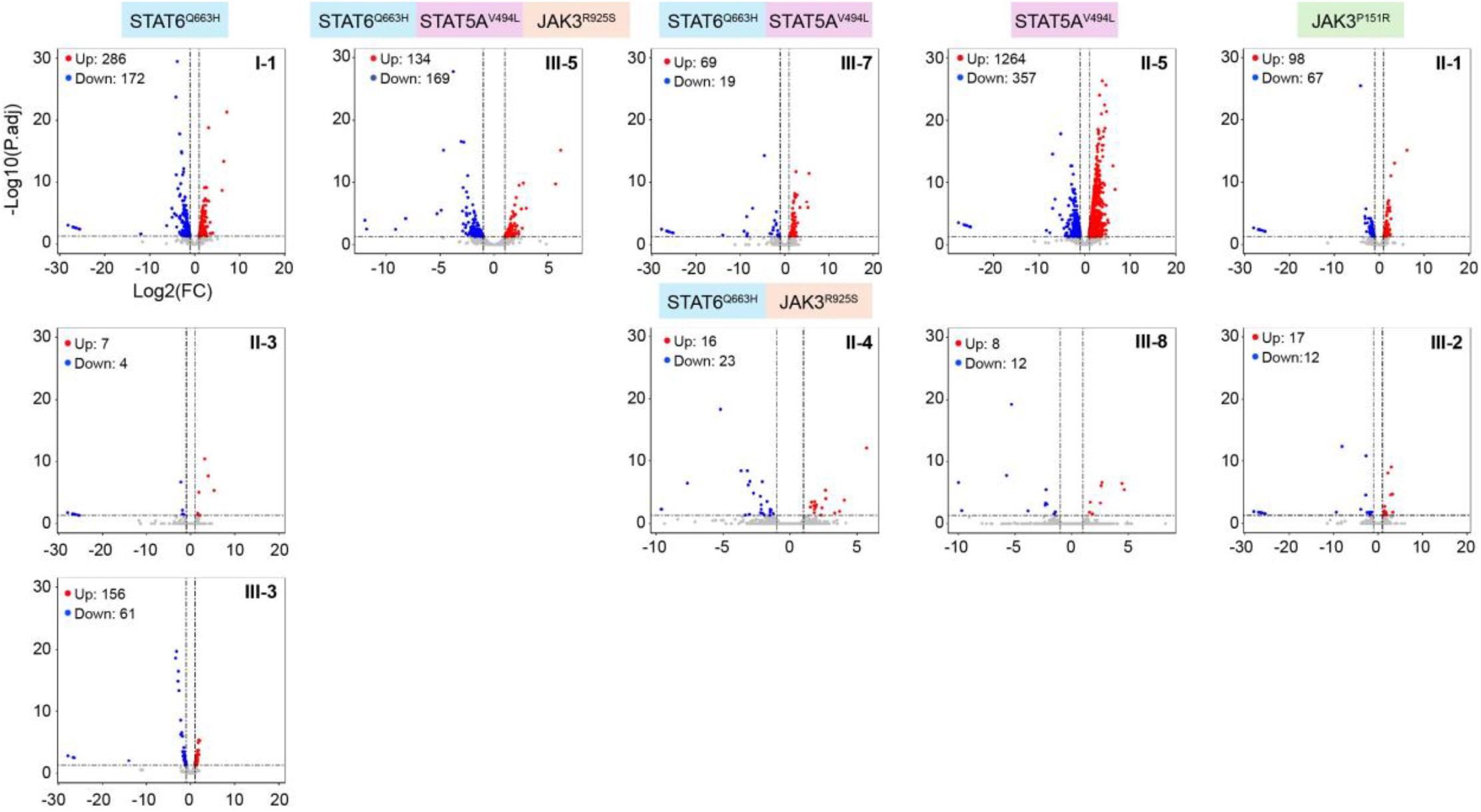
Differentially expressed genes in PBMCs from probands two weeks after SARS-CoV-2 infection. carrying different combinations of JAK/STAT variants.

## Discussion

The biological function of most germline and somatic human JAK/STAT variants remains elusive and the impact of combinatorial SNVs and the possible role of epistasis on physiological responses in humans adds additional layers of complexity. Such biological questions can be studied best in families carrying combinations of different variants. Here we identify four very rare JAK/STAT variants in a Tyrolean family with 17 members and integrate data-driven approaches and RNA transcriptomes with the goal to further elucidate their functions, alone and in combinations. We demonstrate that all four variants are inherited in a Mendelian fashion without any evident clinical or physiological manifestations in the carriers. Most notably, only some carriers of specific variants presented with an altered immune transcriptome suggesting a higher complexity of coinciding JAK/STAT and other immune regulatory SNVs.

With only limited clinical details available of the rare mutations, we employed *in silico* predictions which, however, presented a complex and somewhat contradictory picture, with some tools suggesting potential pathogenicity while others indicate benign effects. While AI-driven AF3 is a very powerful tool to predict protein structures, it provided limited insight into the biological impact of the variants under investigation. The inconsistency among computational predictions highlights the difficulty of depending solely on *in silico* approaches to evaluate how mutations affect complex signaling pathways, emphasizing the importance of experimental confirmation.

Three variants, JAK3^P151R^, JAK3^R925S^ and STAT6^Q633H^ could be suspected as GOF variants because somatic versions are documented in the Catalog of Somatic Mutations in Cancer (COSMIC). Previous literature documented the presence of nine of the eleven known germline STAT6 GOF variants in COSMIC^35-37^. JAK3^R925S^ is a rare variant that has been reported in patients with T-ALL^38^ but by itself it does not transform Ba/F3 cells *in vitro* or stimulate cell proliferation of the MOHITO cell line^39^. While JAK3^R925S^ has been described as a passenger mutation in T-ALL patients, we suggest that it could impact the basal immune transcriptome and virus-induced innate immune responses in the presence of additional JAK/STAT mutation line in proband III-5 who also carries the STAT5A^V494L^ and STAT6^Q633H^ variants. A possible synergy between JAK variants has also been reported in leukemia patients carrying two or more variants^40^. STAT6^Q633H^ is a very rare variant that has not yet been reported in the literature. However, one case has been reported in the COSMIC database and PolyPhen2 reports it as pathogenic. This variant is located at the intersection of the SH2 and TAD domain, in close vicinity of well-known GOF variants, including P643R^36,37^ and the key tyrosine phosphorylation site (Y641). Since only one out of the three individuals in our study carrying the STAT6^Q633H^ variant displayed an elevated basal and SARS-CoV-2 induced innate transcriptome it is likely that additional variants in immune regulators contribute to its function. Several hundred variants were identified in these individuals in other key immune genes, such as TYK2 and interferon receptors, and the complexity of variant interactions remains to be investigated. Like the STAT6 variant, only one of the three individuals carrying the very rare STAT5A^V494L^ displayed an enhanced basal transcriptome, again, suggesting the presence of variants in additional immune regulatory genes.

Our findings highlight the combinatorial complexity of JAK/STAT variants in humans and their likely impact on physiology and pathophysiology. It also underscores the importance of obtaining WGS data in any family study, permitting the identification of cooperating variants in related pathways. This presence of cooperating variants is highlighted by findings that the MPN driver mutation JAK2^V617F^ has been found in large cohorts of healthy elderly individuals^41^. Since mutations in humans do not exist in isolation, their impact can only be gauged in the context of additional mutations in their native setting. Understanding the complexity of epistatic SNP interactions has become a major challenge^42^ in understanding how the co-presence of several variants can modulate physiological and pathophysiological responses. Artificial intelligence driven algorithms will be critical in integrating the data on individual gene variants and their biological and clinical effects.

## Limitations

A limitation of the investigation is that we had identified only one extended family carrying the four JAK/STAT variants.

## Author contributions

LH and LK initially designed the study. As the study progressed, HKL, PAF, TH and OS contributed to further design of the study and data analysis. LK recruited study population and LK and YC collected blood and prepared DNA and RNA. HKL generated and analyzed WES and RNA-seq data. TH and OS conducted AF3 analyses. HKL and LH wrote the manuscript, TH, OS and PAF edited the manuscript and all authors approved the final version. We acknowledge the use of ChatGPT (https://chat.openai.com/) and HHS.AI to provide general information on topics discussed in this work and to clarify context.

## Funding

This research was supported by the Intramural Research Program of the National Institute of Diabetes and Digestive and Kidney Diseases (NIDDK) within the National Institutes of Health (NIH). The contributions of the NIH author(s) were made as part of their official duties as NIH federal employees, are in compliance with agency policy requirements, and are considered Works of the United States Government. However, the findings and conclusions presented in this paper are those of the author(s) and do not necessarily reflect the views of the NIH or the U.S. Department of Health and Human Services. Additional funding was obtained from the Research Council of Finland, Sigrid Juselius Foundation, Finnish Cancer Foundation, Tampere Tuberculosis Foundation, Competitive Research Funding of the Tampere University Hospital – Fimlab.

## Acknowledgements

This research was supported by the Intramural Research Programs (IRPs) of National Institute of Diabetes and Digestive and Kidney Diseases (NIDDK) within the National Institutes of Health (NIH). The contributions of the NIH author(s) are considered Works of the United States Government. The findings and conclusions presented in this paper are those of the author(s) and do not necessarily reflect the views of the NIH or the U.S. Department of Health and Human Services.

We thank the NIH Intramural Sequencing Center, NISC (https://www.nisc.nih.gov/contact.htm) for NGS and RNA-seq. This work utilized the computational resources of the NIH HPC Biowulf cluster (http://hpc.nih.gov).

---

**Supplementary Table 1.**
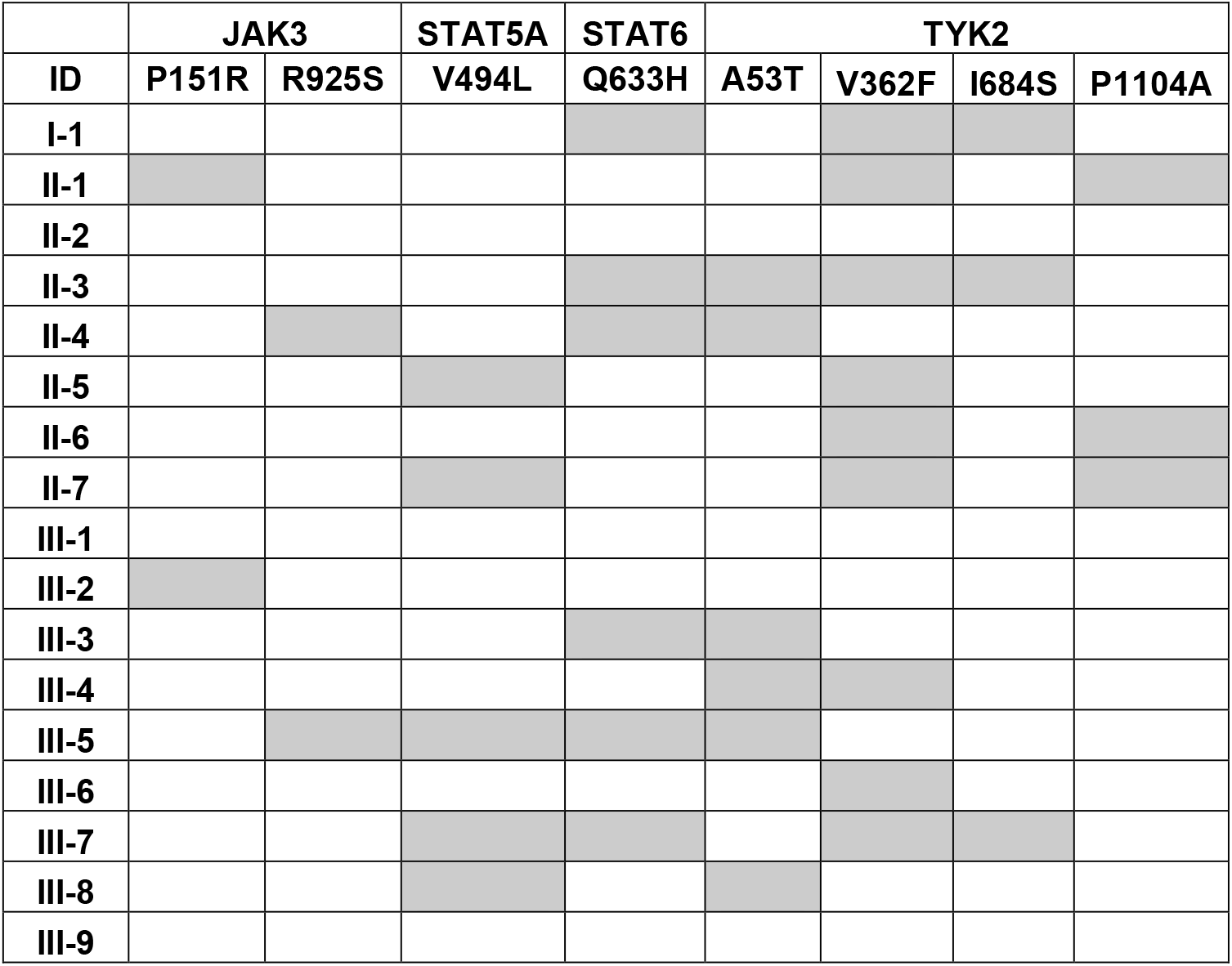
Combinatorial occurrence of JAK/STAT variants in study cohort.

**Supplementary Table 2. List of genes significantly regulated in the individual compared with the control group**.

**Supplementary Table 3. List of genes significantly regulated in the individual compared with the control group at Day14 after Omicron infection**.

## Notes

### Competing Interest Statement

The authors have declared no competing interest.

